# Induction of DISE in ovarian cancer cells *in vivo*

**DOI:** 10.1101/141945

**Authors:** Andrea E. Murmann, Kaylin M. McMahon, Ashley Haluck-Kangas, Nandini Ravindran, Monal Patel, Calvin Law, Sonia Brockway, Jian-Jun Wei, C. Shad Thaxton, Marcus E. Peter

## Abstract

The death receptor CD95/Fas can be activated by immune cells to kill cancer cells. shRNAs and siRNAs derived from CD95 or CD95 ligand (CD95L) are highly toxic to most cancer cells. We recently found that these sh/siRNAs kill cancer cells in the absence of the target by targeting the 3’UTRs of critical survival genes through canonical RNAi. We have named this unique form of off-target effect DISE (for death induced by survival gene elimination). DISE preferentially kills transformed cells and cancer stem cells. We demonstrate that DISE induction occurs in cancer cells *in vivo* after introducing a lentiviral CD95L derived shRNA (shL3) into HeyA8 ovarian cancer cells grown as i.p. xenografts in mice, when compared to a scrambled shRNA. To demonstrate the possibility of therapeutically inducing DISE, we coupled siRNAs to templated lipoprotein nano particles (TLP). *In vitro,* TLPs loaded with a CD95L derived siRNA (siL3) selectively silenced a biosensor comprised of Venus and CD95L ORF and killed ovarian cancer cells. *In vivo,* two siRNA-TLPs (siL2-TLP and siL3-TLP) reduced tumor growth similarly as observed for cells expressing the shL3 vector. These data suggest that it is possible to kill ovarian cancer cells *in vivo* via DISE induction using siRNA-TLPs.

## INTRODUCTION

CD95/Fas is a death receptor that, together with its ligand CD95L, regulates immune homeostasis [1, 2]. Immune cells such as cytotoxic killer and natural killer (NK) cells use CD95L to kill virus infected and cancer cells [3]. However, we, and others, reported that CD95 and CD95L have multiple tumor promoting activities [4–6] and tissue specific deletion of CD95 in the liver or the ovaries of mice strongly reduced or prevented tumor formation in these tissues [7, 8]. We found that >80% of 22 different siRNAs, DsiRNAs or shRNAs targeting either CD95 or CD95L killed cancer cells [8, 9] through a process we had coined DICE (for death induced by CD95/CD95L elimination). DICE is independent of caspase-8, RIPK1, MLKL, and p53, is not inhibited by Bcl-x_L_ expression, and it preferentially affects cancer cells [8]. It is characterized by an increase in cell size and production of mitochondrial ROS, which is followed by DNA damage. It resembles a necrotic form of mitotic catastrophe. No single drug was found to completely block this form of cell death, and DICE could also not be blocked by the knockdown of any single gene, making it a promising new way to kill cancer cells [8]. More recently, we reported that DICE preferentially affects cancer stem cells [10] suggesting a physiological role of DICE in targeting neoplastically transformed cells.

Surprisingly, we recently discovered that DICE works even in the complete absence of CD95 and CD95L [9]. DICE is, therefore, a highly selective form of an RNAi off target effect (OTE). We found that CD95 and CD95L mRNAs contain dozens of sequences that target a network of genes found to be critical for the survival of cancer cells and that are often upregulated in cancer [9]. These sequences, when introduced into cancer cells in the form of transfected siRNAs or lentiviral shRNAs, act through canonical RNAi, targeting the survival genes through short seed matches in their 3’UTRs. The cancer cells likely die due to the loss of multiple survival genes. We have therefore called this form of cell death DISE (for death induced by survival gene elimination) [9]. The complex nature of this toxicity may explain why cancer cells have difficulties developing resistance to DISE.

We have now explored how inducing DISE may be a novel form of cancer therapy. We first demonstrated that DISE induction works in an *in vivo* model whereby mice harbor xenografted ovarian cancer cells expressing a CD95L derived shRNA (shL3). We then delivered two siRNAs derived from CD95L (siL2 and siL3) to mice with ovarian cancer xenografts using a nanoparticle platform demonstrated previously to deliver siRNA *in vivo* [11, 12]. Templated lipoprotein particles (TLP) stabilize siRNA and are dependent upon SR-B1 expression for efficient siRNA delivery. The TLPs were delivered i.p., taken up by the tumor cells, and acted through canonical RNAi, substantially reducing tumor growth.

The *in vivo* study was terminated once the control treated mice showed signs of discomfort due to large tumors and/or ascites formation. The remaining tumor cells from the nanoparticle delivered siL3 group were resected, cultured, and transfected with siL3 in culture using a commercially available cationic lipid-based transfection reagent in order to determine if the cells became insensitive to DISE. In this context, the tumor cells were still fully susceptible to DISE, suggesting that treatment optimization (i.e. dose and time) may allow for full eradication of tumor growth. Collectively, these data demonstrate that DISE induction is a promising new approach for treating cancer.

## RESULTS

### Induction of DISE using shRNAs *in vivo*

In multiple cancer cells tested *in vitro*, including ovarian cancer cell lines (Supplementary Figure S1A), DISE induction is a potent mechanism for cancer cell treatment [8]. To test whether DISE is a viable mechanism for *in vivo* treatment, we chose an orthotopic mouse model of ovarian cancer. We selected the highly active shRNA shL3, which we have previously demonstrated kills cancer cells by targeting survival genes [9]. *In vitro*, cells were selected with puromycin after lentiviral infection. However, because puromycin selection could not be used *in vivo*, prior to *in vivo* experiments we tested the efficiency of shRNA virus infection and DISE induction in HeyA8 Venus-siL3-pfuL2T cells with and without puromycin selection (Figure 1A). Cell growth was reduced in cells treated using a MOI of 5 (without puromycin treatment) and was comparable to cells selected by treatment with puromycin (after infection with a MOI of 3). The *in vitro* data demonstrate that DISE induction occurs roughly 2-3 days after introducing the shRNA [8]. To determine the effects of DISE *in vivo*, we injected NSG mice i.p. with cells infected with either pLKO-shScr or pLKO-shL3 virus without puromycin treatment (Figure 1B, left panel) or treated for one day with puromycin before injection (Figure 1B, right panel). Tumor growth was monitored over two weeks and tumor cells expressing shL3 grew slower when compared to tumors expressing shScr. Upon histological inspection of the tumors by a pathologist, 13 days after cancer cell injection, shScr control treated tumors showed large areas of central necrosis in larger tumors, presumably caused by hypoxia (Figure 1C-a, b, top panel), and demonstrated signs of tumor cells invading the omentum with no signs of necrosis (Figure 1C-c, top panel). The shL3 expressing tumors showed signs of necrosis in smaller tumors than seen in the shScr expressing tumor cells (Figure 1C-d). Clear signs of tumor regression were seen with small areas of residual tumors. Our data demonstrate that cancer cells, when grown *in vivo*, are susceptible to DISE, confirming that DISE induction may be a potent therapy for cancer.

**Figure 1:**
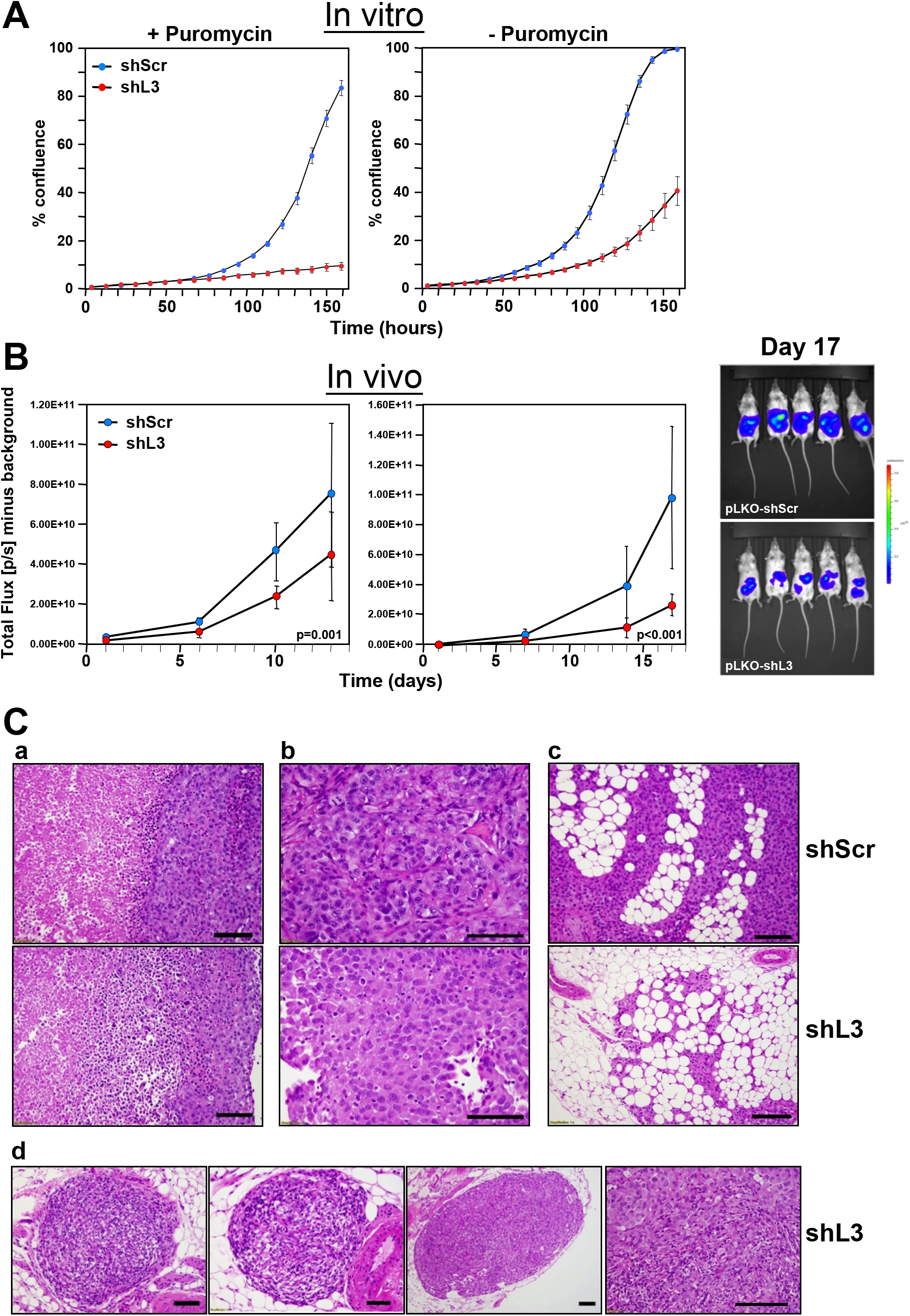
Expression of a CD95L derived shRNA causes induction of DISE *in vitro* and *in vivo*. **A:** Percent growth change over time of HeyA8 Venus-siL3-pfuL2T cells infected with either shScr or shL3 pLKO lentiviruses (MOI = 5) (with and without puromycin selection) at 250 cells/well. **B:** Small animal imaging of HeyA8 pfuL2T cells infected with either shScr or shL3 pLKO lentiviruses (MOI = 5) after i.p. injection into NSG mice (10 mice per group, 10^6^ cells/mouse). *Left:* mice injected with cells infected with virus without puromycin selection; *Center:* Mice injected with HeyA8 cells infected with shRNAs and selected with puromycin for 24 hours. *Right:* Bioluminescence image of 5 mice 17 days after i.p. injection with HeyA8 cells infected with either shScr or shL3 virus. Two-way ANOVA was performed for pairwise comparisons of total flux over time between shScr and shL3 expressing cells. **C:** H&E staining of representative tumors isolated from mice carrying HeyA8-shScr (a,b,c, top row) and HeyA8-shL3 tumors (a,b,c, bottom row and d). a, in shScr treated tumors, tumor mass showed two zones of viable (right) and necrotic (left) tumor regions with sharply demarked boundary. The viable tumor cells were cohesive with dense basophilic and pale cytoplasm. In shL3 treated tumor, a zone of dying tumor cells sites were seen in between viable and necrotic zones. This zone had tumor cells that were loosely cohesive with mixed dying, dead and viable cells. b, Close view of tumor cells revealed the different cytologic features. In shScr treated tumors, cells were more cohesive with a solid growth pattern with centrally located large and high grade nuclei. In shL3 treated tumors, cells were loosely cohesive with eccentrically located nuclei and eosinophilic and hyaline cytoplasm. These findings suggest early degenerative or regressing changes. c, Tumor infiltrating into fat had minimal or no tumor cell necrosis. In shScr treated tumors, tumor mass in fat had large and high tumor volume (top panel). In shL3, infiltrating tumor cells were much smaller in size and volume and areas of regression change were seen (bottom panel). d, Tumor regression could be frequently seen in shL3 treated tumors, characterized by well demarked tumor nodules (left three images) with peripheral rim of viable tumor cells (right panel) and central regression of tumor bed which was replaced by histiocytes, lymphocytes and fibrotic stromal cells.

To induce shRNA expression after injection of tumor cells, we used the Tet inducible pTIP vector [8]. HeyA8-pFuL2G cells were stably infected with either pTIP-shScr or pTIP-shL3 in the presence and absence of puromycin selection and then treated with Doxycycline (Dox) to induce shRNA expression. shL3 expression significantly slowed down growth of cells compared to shScr (Figure 2A). Puromycin treatment did not have an effect on cell growth in the absence of Dox. HeyA8 cells treated with Dox to induce shL3 showed little to no growth and most cells demonstrated cell death when compared to shScr cells (Movies S1 and S2). The pTIP-shScr and pTIP-shL3 cells were injected i.p. into NSG mice and one day after tumor injection half the mice were given Dox in their drinking water. Small animal imaging showed that tumor growth was significantly reduced in mice injected with cells expressing pTIP-shL3 that received Dox (Figure 2B, left panel). The level of growth reduction was more pronounced when the tumors were excised and weighed (Figure 2B, right panel).

**Figure 2:**
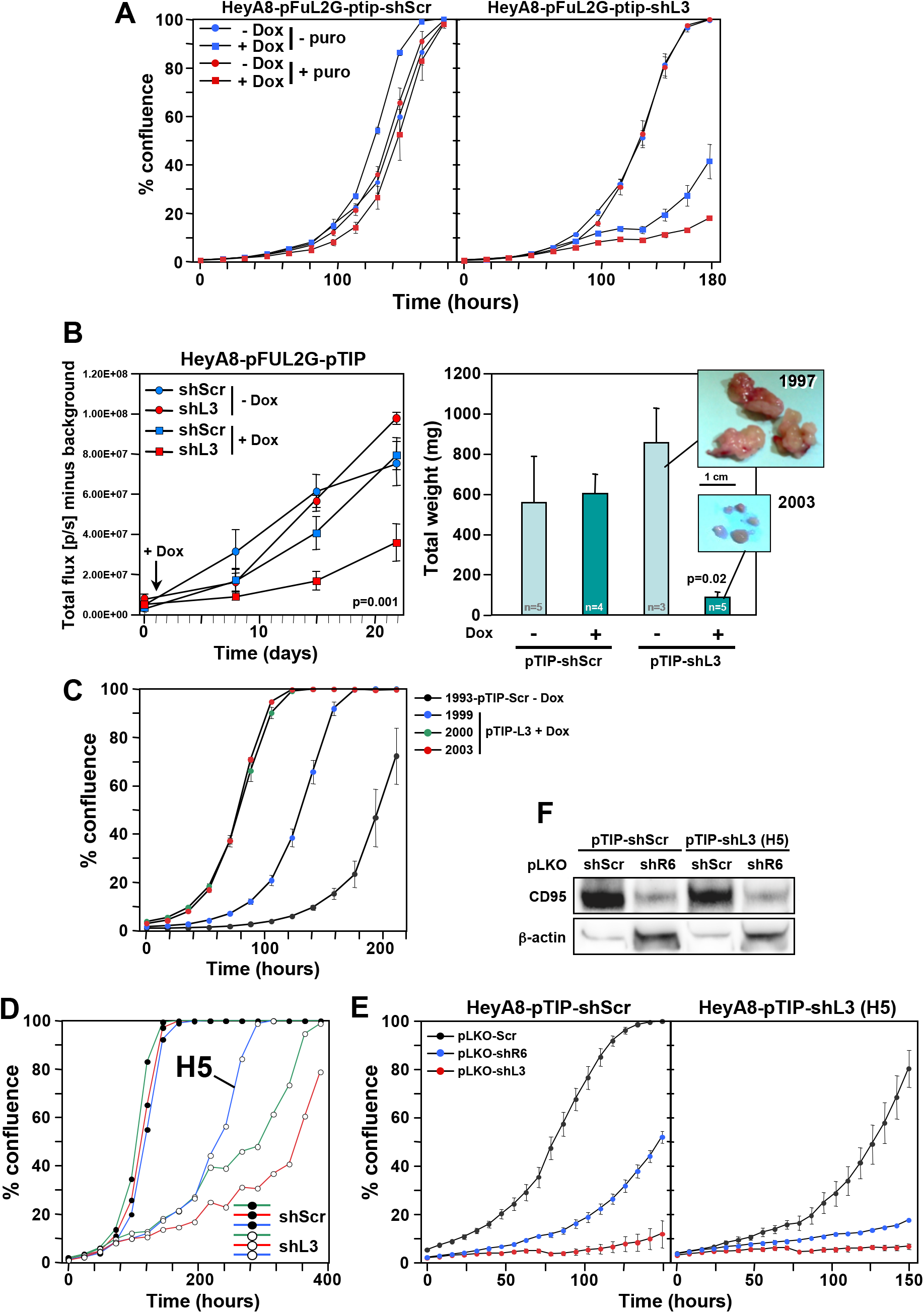
HeyA8 cells regress *in vivo* after inducible expression of shL3 and become resistant to the inducible vector but not to DISE induction. **A:** Percent growth change over time of HeyA8-pFul2G cells (plated at 250 cells per 96 well) expressing either pTIP-shScr or pTIP-shL3 cultured with or without Dox. **B:** Small animal imaging of HeyA8-pFul2G cells expressing either pTIP-shScr or pTIP-shL3 after i.p. injection into NSG mice (10^6^ cells/mouse). *Left:* Tumor growth over time. The day the mice were given Dox containing drinking water is labeled with an arrow. ANOVA was performed for pairwise comparisons of total flux over time between shScr and shL3 expressing cells. *Right:* Tumor weight in each treatment group 22 days after tumor cell injection (pictures of the tumors of two representative mice are shown). P-value was calculated using Student’s ttest. **C:** Change in confluence over time of four tumors isolated from 4 mice in B treated as indicated all in the presence of Dox. **D:** Change in confluency of HeyA8 cells expressing either pTIP-shScr or pTIP-shL3 in the presence of Dox. Confluency of three wells each is shown. The Dox resistant clone H5 was chosen for further analysis. **E:** Change in confluency of Hey8-pFul2G cells or the H5 clone over time in the presence of Dox after infection with either pLKO-shScr, pLKO-shL3 or pLKO-shR6 (MOI=6). **F:** Western blot analysis for CD95 of the cells in E infected with either pLKO-shScr or pLKO-shL3.

Because there was remaining pTIP-shL3 tumor at the end of the study, we wanted to determine if the cells were no longer responsive to shL3 due to a mutation, had become resistant to DISE, had reduced function of the RNAi machinery, or no longer responded to Dox. Therefore, we isolated cells from tumors of four mice and cultured them in puromycin for 2 days to eliminate any host cells. Three resected tumors were from pTIP-shL3 mice that had received Dox, and the fourth tumor was from a mouse carrying a pTIP-shScr expressing tumor. All four cultured tumors expressed roughly the same amount of GFP documenting that a substantial number of cells in each culture were HeyA8 cells with the pTIP vector (Supplementary Figure S2). When these cells were plated in the presence of Dox all four of them grew *in vitro* (Figure 2C). To determine if mutations present in the cell culture were responsible for the apparent resistance to toxic activity of shL3, we cultured HeyA8-pTIP-shScr and HeyA8-pTIP-shL3 cells *in vitro* in the presence of Dox for an extended period of time. Interestingly, every clone plated grew out (Figure 2D and data not shown, Movie S3, well H5 and Movie S4, control). However, it was unlikely that rare mutations occurred in every well that contributed to the clone growth, because only 250 cells were plated per well. To determine if cells had become resistant to the DISE inducing activity of shL3 we introduced the shL3 shRNA in a different form into one of the resistant clones (H5) using the pLKO vector in the presence of Dox and puromycin (Figure 2E). While the H5 cells were impaired in their growth compared to cells isolated from a well with cells expressing shScr, they were not resistant to DISE induced by either shR6 (a shRNA targeting CD95) or shL3 shRNA (Figure 2E, right panel). These data suggest that RNAi was still fully active in the resistant clone. This was also confirmed by the demonstration that shR6 knocked down CD95 as efficiently in the H5 clone as in the control cells (Figure 2F). Most important however, was the observation that the H5 clone which was resistant to Dox induction of shL3 driven from the pTIP vector was still fully susceptible to the very same shRNA when induced using a different vector. The *in vitro* data suggest that tumor cells did not develop resistance to DISE but rather to the vector or to Dox used to induce the shRNA. Similar data were also obtained *in vitro* and *in vivo* when the pTIP-shL3 vector was used in a breast cancer mouse model in which we injected MB-MDA-231 cells expressing pTIP-shScr or pTIP-shL3 into the fat pad and cultured the cells ex vivo (data not shown). This suggests that the resistance that developed is not specific to HeyA8 cells but could be a more fundamental effect seen with the pTIP vector.

### siL3 kills ovarian cancer cells by targeting a network of survival genes

These data suggested that it might be possible to induce DISE *in vivo* by delivering siRNAs to tumor cells. We recently showed that the majority of the tested shRNAs, siRNAs and DsiRNAs that are derived from CD95L induced DISE [9]. We selected two toxic siRNAs, siL2 and siL3, derived from CD95L. Both were active in silencing their cognate target [8] and in reducing growth of HeyA8 cells (Figure 3A). To monitor delivery and RNAi activity of siL3 *in vivo* we developed a biosensor plasmid that carried a CD95L mini gene of 50 nt with the center comprised of the 20 nt sequence that is targeted by siL3. This minigene was linked to a Venus fluorophore (Figure 3B). We generated a mutant form of the sensor whereby the siL3 target site was mutated in 6 positions. HeyA8 cells were generated either expressing the siL3WT or the siL3MUT sensor. These cells were transfected with a nontargeting siRNA (siScr), siL3 or siL3MUT and, after 2 days, cells were analyzed by flow cytometry (Figure 3C). The green fluorescence in the cells carrying the two sensor constructs was only reduced upon transfection with the siRNA that was completely complementary to the sequence in the sensor. This established the sensor system as a sensitive tool to detect the activity of the CD95L targeting siL3. Finally, we monitored the change in fluorescence, cell morphology and confluency (as a surrogate marker of cell viability) over time upon transfection with the different siRNAs (Figure 3D and Movies S5-S8). When HeyA8 cells expressing the siL3WT Venus sensor were transfected with siL3MUT they remained green and became confluent after 6 days (Movie S5). The same cells transfected with siL3 lost green fluorescence after about 2 days and never reached confluency due to DISE induction (Movie S6). In contrast, when HeyA8 cells expressing the siL3MUT sensor were transfected with the complementary siL3MUT siRNA they lost green fluorescence after 2 days but became confluent (Movie S7), when transfected with siL3 they remained green but did not reach confluency due to DISE induction (Movie S8). These experiments confirmed that siL3 induces DISE in a sequence specific manner.

**Figure 3:**
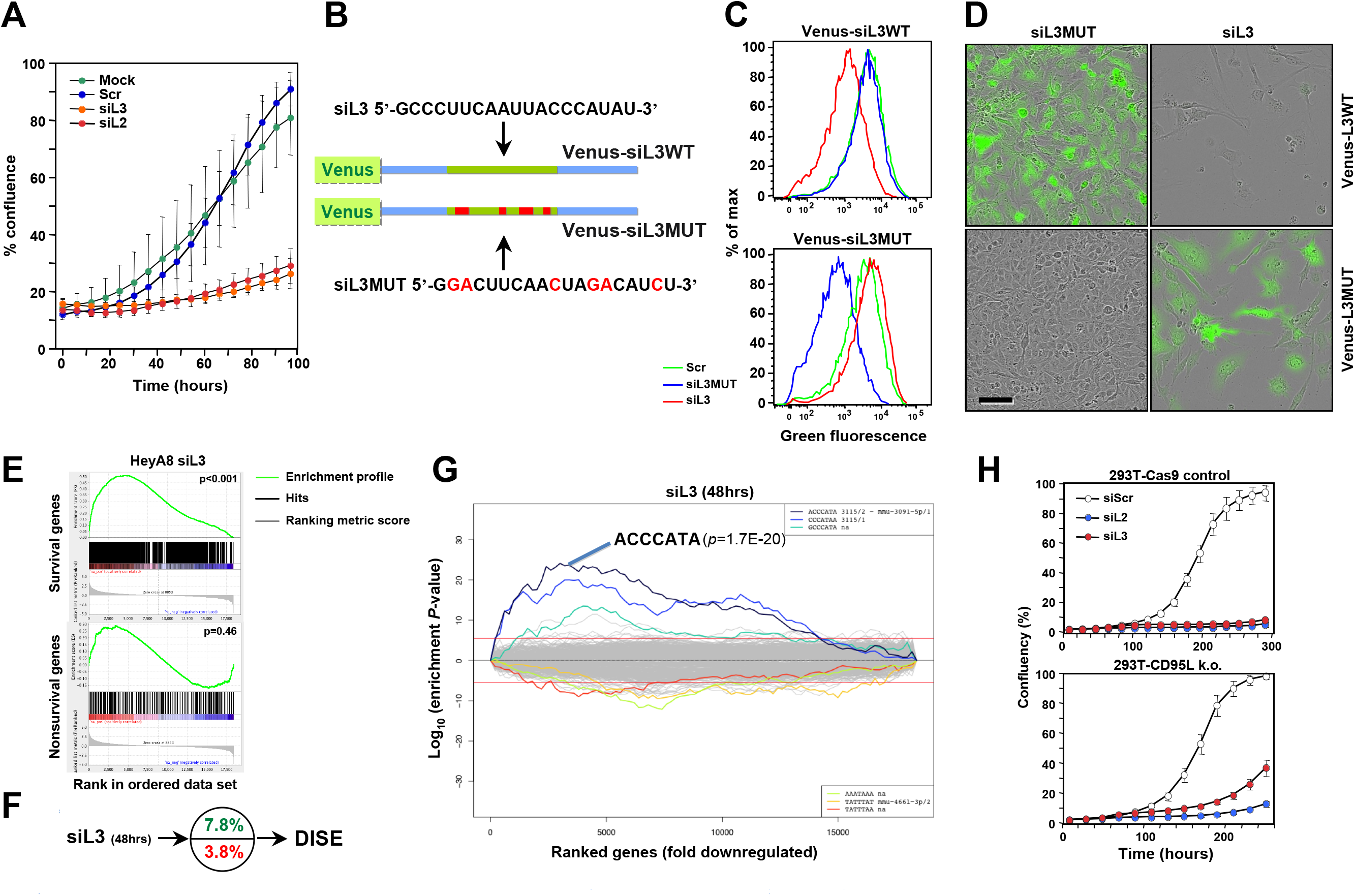
CD95L derived siL2 and siL3 knockdown CD95L and induce DISE by targeting critical survival genes. **A:** Specific targeting of a minisensor containing Venus fused to a CD95L minigene comprised of the 50 nt surrounding the siL3 target site in CD95L. Cells were either mock transfected (Lipofectamine only) or transfected with either 5 nM siScr, siL2 or siL3. Shown is change in percent confluency over time when cells were plated at 3000 cells per well in a 48-well plate 24 hours after transfection. **B:** Schematic showing the design of the wt Venus-siL3 sensor and a sensor carrying a mutated siL3 targeted site. Sequences of targeting siRNAs are given. **C:** FACS analysis of HeyA8 cells stably expressing either the wt or the mutant Venus-siL3 sensor (see B) 24 after transfection with 10 nM of either siScr, siL3WT or siL3MUT. **D:** Composite images of phase contrast and green fluorescence of HeyA8 cells expressing either Venus-siL3WT or Venus-siL3MUT sensor 6 days after transfection with 10 nM of either siL3 or siL3MUT. Scale bar = 150 μm. **E:** Enrichment of the siL3 seed match in the 3’UTRs of downregulated survival genes. **F:** Preferential downregulation of survival genes in cells transfected with siL3 as in E. **G:** SYLAMER analysis of HeyA8 cells 50 hrs after transfection with 25 nM siL3. **H:** Change in confluency over time of a mix of three 293T-Cas9 clones compared to a mix of two complete 293T CD95L deletion clones after transfection with 25 nM siScr, siL2 or siL3 (all from Dharmacon, IDT oligonucleotides gave very similar results (data not shown)).

We recently showed that a number of siRNAs and shRNAs derived from either CD95 or CD95L are toxic to cancer cells because they target a network of genes that are critical for the survival of the cells. We demonstrated that siL3 can kill HeyA8 cells even after homozygously deleting the siL3 targeted site in HeyA8 cells [9]. To test whether the naked siL3 oligonucleotides we planned to couple to nanoparticles killed cancer cells by targeting survival genes we subjected HeyA8 cells with no detectable expression of CD95L (data not shown) and transfected with unmodified siScr or siL3 and then performed RNA Seq analysis. The most downregulated genes in the siL3 treated cells when compared to the cells treated with control siScr were enriched in genes recently described in a genome wide CRISPR based lethality screen [13] as survival genes (Figure 3E) but not enriched in nonsurvival genes. In fact, survival genes were about two times more likely targeted than nonsurvival genes (Figure 3F). Performing a Sylamer analysis which identifies seed matches of si- and miRNAs that are enriched in the 3’UTRs of a ranked list of genes, confirmed an enrichment of the siL3 seed match in the 3’UTRs of the downregulated genes (Figure 3G). The data in Figures 3E–3G were very similar to the ones obtained for cells treated with chemically modified siL3 suggesting that the naked siRNA had comparable activities [9]. To test whether an independent toxic siRNA, siL2, derived from CD95L also induced DISE we generated 293T cells lacking the entire CD95L locus using CRISPR/Cas9 gene editing (Supplementary Figure S3). Transfection of these 293T CD95L k.o. cells with either siL3 and siL2 resulted in substantial cell growth inhibition (a surrogate marker of cell death) (Figure 3H) suggesting that siL2, just like siL3, is a toxic siRNA derived from the CD95L gene that kills cancer cells by DISE.

### Induction of DISE in cancer cells using TLP nanoparticles *in vitro* and *in vivo*

We have recently developed a new form of siRNA delivery using TLP nanoparticles [11]. We described that uptake of the TLPs was dependent on expression of the scavenger receptor SR-B1 [11]. To test which of the cancer cell lines expressed SR-B1, we subjected a number of solid and blood cancer cell lines to Western blot analysis (Figure 4A). SR-B1 was expressed in all tested cancer cell lines at varying levels, except for Jurkat cells. We then tested uptake of TLPs coupled with a Cy5 labeled siL3 oligonucleotide (Figure 4B). Both fluorescence microscopy and FACS analysis demonstrated efficient uptake of the labeled particles in 17 different cancer cell lines. Particles were taken up within 1 hour and when excess particles were washed away detection of labeled siL3 slowly dissipated (Figure 4C). We next tested whether uptake of siL3 loaded TLPs resulted in reduction of green fluorescence and induction of DISE in HeyA8-Venus-siL3 cells (Figure 4D). The siL3-TLP, but not the siL3MUT-TLP (see Figure 3B) or empty TLPs, had a strong effect on green fluorescence (Figure 4D and 4E, F). Incubation of these cells with the siL3-TLP also induced DISE (Figure 4D and F). This was found for the p53 mutant ovarian cancer cell line OVCAR3 and for another epithelial ovarian cancer cell line OVCAR4 (Figure S1B).

**Figure 4:**
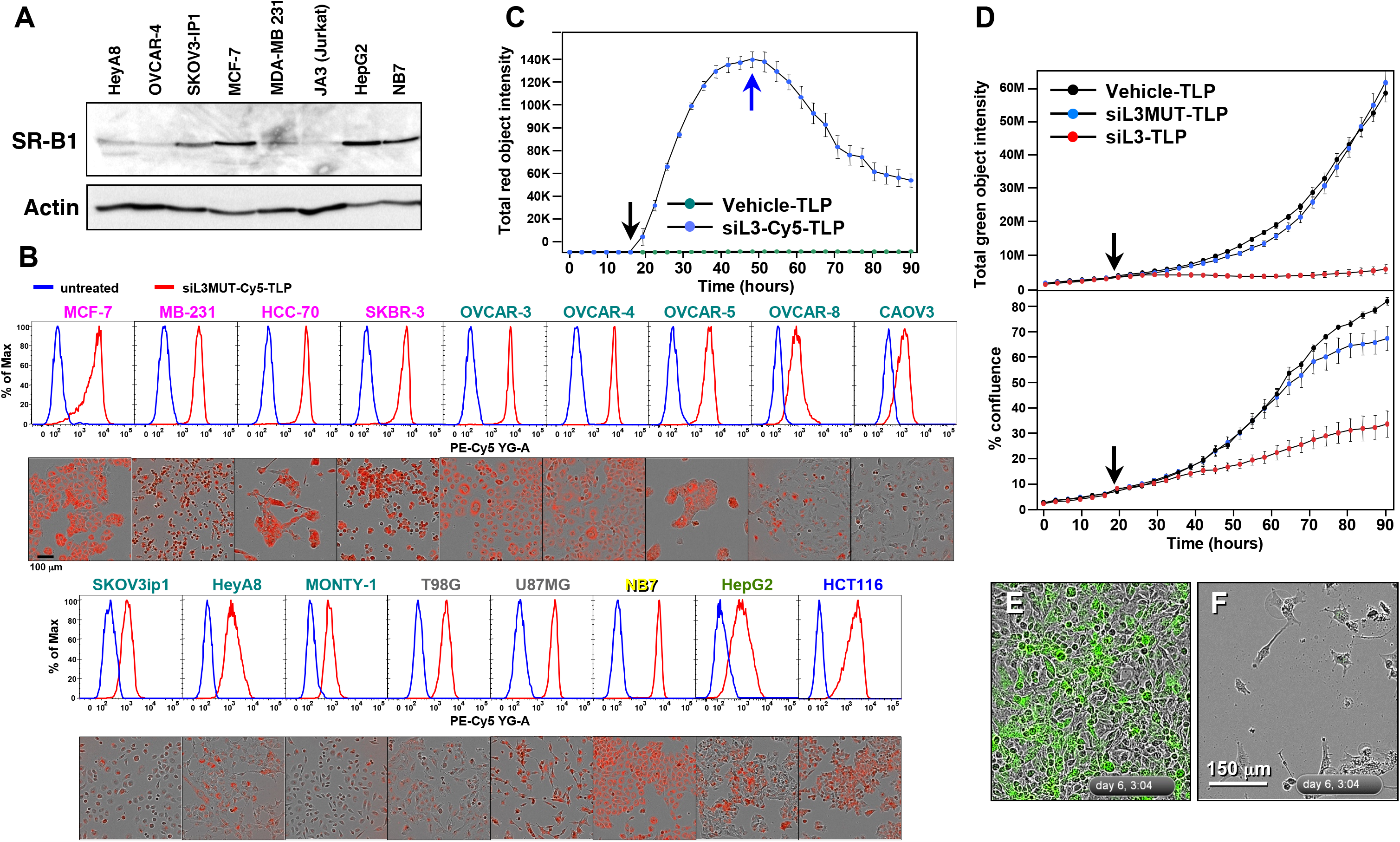
siL3-TLP uptake and DISE induction. **A:** Western blot analysis for SR-B1 in different cancer cell lines. **B:** Uptake of siL3MUT-Cy5-TLP by 17 different cancer cell lines representing 6 different cancers, detected by flow cytometry (top row) and phase contrast/red fluorescence (bottom). **C:** Change of red (Cy5) fluorescence (red object count) over time of HeyA8 cells treated with TLPs. Black arrow, particles were added; Blue arrow, cells were washed and particles removed. **D:** HeyA8 cells expressing the Venus-siL3 sensor were incubated for 90 hours with 8 nM (RNA concentration) siL3-TLP, siL3MUT-TLP, or Vehicle-TLPs. Cells were analyzed in an IncuCyte Zoom. *Top:* Total (Venus) green fluorescence. *Bottom:* Confluency. **E, F:** Images (merged phase contrast and green fluorescence) of the cells after a 147 hours incubation with either Vehicle-TLP (E) or siL3-TLP (F).

To test whether siRNA-TLPs could be used to induce DISE *in vivo*, HeyA8-Venus-siL3 cells also expressing a Tomato red luciferase construct were injected i.p. into NSG mice and a day after injection mice were subjected to IVIS analysis. Based on the bioluminescence signal they were sorted into three groups (10 mice per group) so that each group had a similar signal distribution. Mice were injected 5 times over 9 days with water, siScr-TLP or siL3-TLP (Figure 5A). Tumors of a small group of additional mice were analyzed by flow cytometry 10 days after tumor injection (after 4 injections with TLCs). The red/green ratio as a measure of targeting the Venus siL3 sensor was determined (Figure 5B). Together with a reduction in green fluorescence in the tumors treated with siL3-TLP when compared to tumors treated with siScr-TLP (Figure 5C) provided evidence that the TLPs had entered the tumor tissue and were actively inducing RNAi in the case of siL3-TLP. We also noticed an increase in the side scatter of red tumor cells taken from three tumors from a mouse treated with siL3-TLP when compared to cells of three tumors from a mouse treated with siScr-TLP (Figure 5D, left panel), a phenomenon also observed *in vitro* in HeyA8 cells after transfection with siL3 (Figure 5D, right panel, and data not shown). The most likely reason for this observation is an increase in granularity of the cells caused by the appearance of massive stress granules caused by DISE induction in these cells [8]. Finally, data show a significant reduction in IVIS signal from the tumor cells at both the second and third measurement (Figure 5E) when treated with the siL3-TLP, but not when treated with water or siScr-TLP. The data suggest that it is possible to deliver a DISE inducing siRNA to tumor cells *in vivo*.

**Figure 5:**
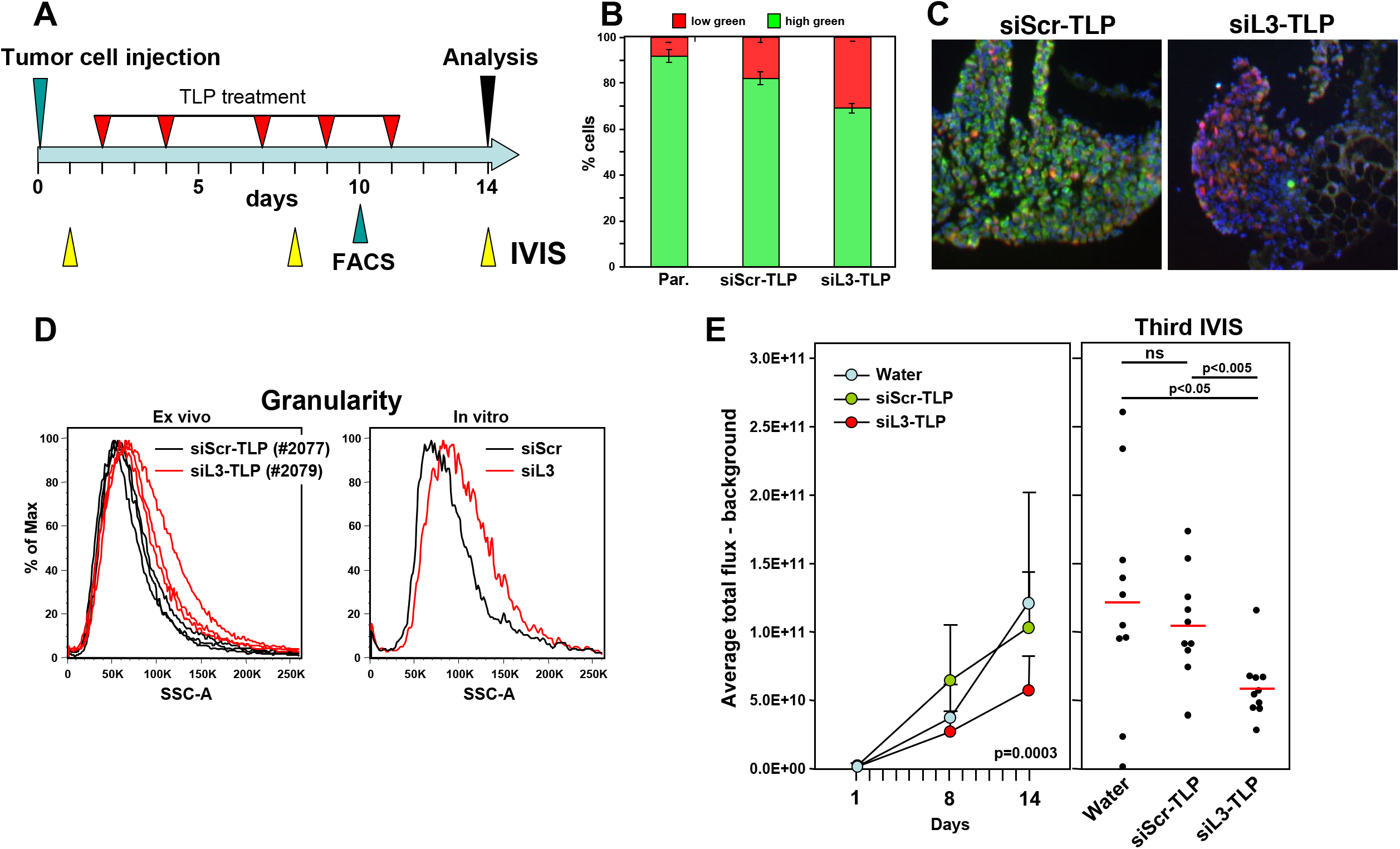
Induction of DISE *in vivo*. **A:** Treatment scheme. **B:** Red/green ratio of tumor cells isolated from three mice each treated with either siScr-TLP and siL3-TLP compared to parental HeyA8-Venus-siL3-pFul2T cells. **C:** Immunofluorescence images of representative tumors from mice treated with either siScr-TLP or siL3-TLP. Size bar = 300 μm. **D:** *Left:* Change in sideward scatter (granularity) of cells isolated from three tumors from two mice treated with either siScr-TLP or siL3-TLP. *Right:* Change in granularity in HeyA8 cells in which DISE was induced by transfection of siL3 (2’O-methylated, Dharmacon) compared to matching siScr. **E:** Small animal imaging of 5 x 10^5^ HeyA8-pFul2G cells injected i.p. into NGS mice treated with either water, siScr-TLPs or siL3-TLPs. *Left:* Tumor growth over time. ANOVA was performed for pairwise comparisons of average flux over time between siScr and siL3 treated cells. *Right:* Bioluminescence signal in individual mice at the third IVIS (14 days after tumor injection) treated as indicated following the treatment protocol outlined in A. P-values were calculated using Student’s ttest.

To test whether a longer treatment and another DISE inducing siRNA would promote a more pronounced tumor growth reduction, we treated mice (n=10/group) with siScr-TLP, siL3-TLP and siL2-TLP for three weeks (Figure 6A). To be able to detect CD95L specific RNAi by the two siRNAs we used HeyA8 cells expressing a Venus sensor that contained the entire CD95L open reading frame [9]. In addition, to be able to distinguish tumor from host tissues in the mice we also stably expressed a red fluorophore localized to the nucleus in the tumor cells. Mice were i. p. injected with 5 x 10^5^ cells and treated with TLP nanoparticles as indicated (Figure 6A). Both siL2-TLP and siL3-TLP substantially reduced tumor growth (Fig 6B). Mice had to be sacrificed at 23 days due to the aggressive growth of HeyA8 cells in siScr-TLP treated mice. To determine whether particles were taken up by the tumor cells and whether the siRNAs acted through RNAi we isolated and froze tumor tissue from three mice in each treatment group and determined by immunofluorescence microscopy the level of reduction in green fluorescence. Overall, reduction in green fluorescence was detectable throughout the tumors of mice treated with the siL2-TLP or siL3-TLP (Figure 6C). While the nanoparticles accumulated on the tumors (Supplementary Figure S4A) no particles were obvious on the livers or the kidneys of the treated mice and no signs of toxicity were detected (Supplementary Figure S4B). This was corroborated by the analysis of the serum of treated mice. No elevation in liver enzymes was measured (Supplementary Figure S4C). Due to the mechanism of DISE induction we found cross-reactivity of human derived sh- and siRNAs to targeted survival genes in mouse cells (data not shown) and transfection of siL3 into mouse ovarian cancer cell line ID8 slowed down their growth (Supplementary Figure S4D). These data suggest that siL3-TLP was not toxic to the mice but reduced tumor growth by inducing DISE in the tumor cells.

**Figure 6:**
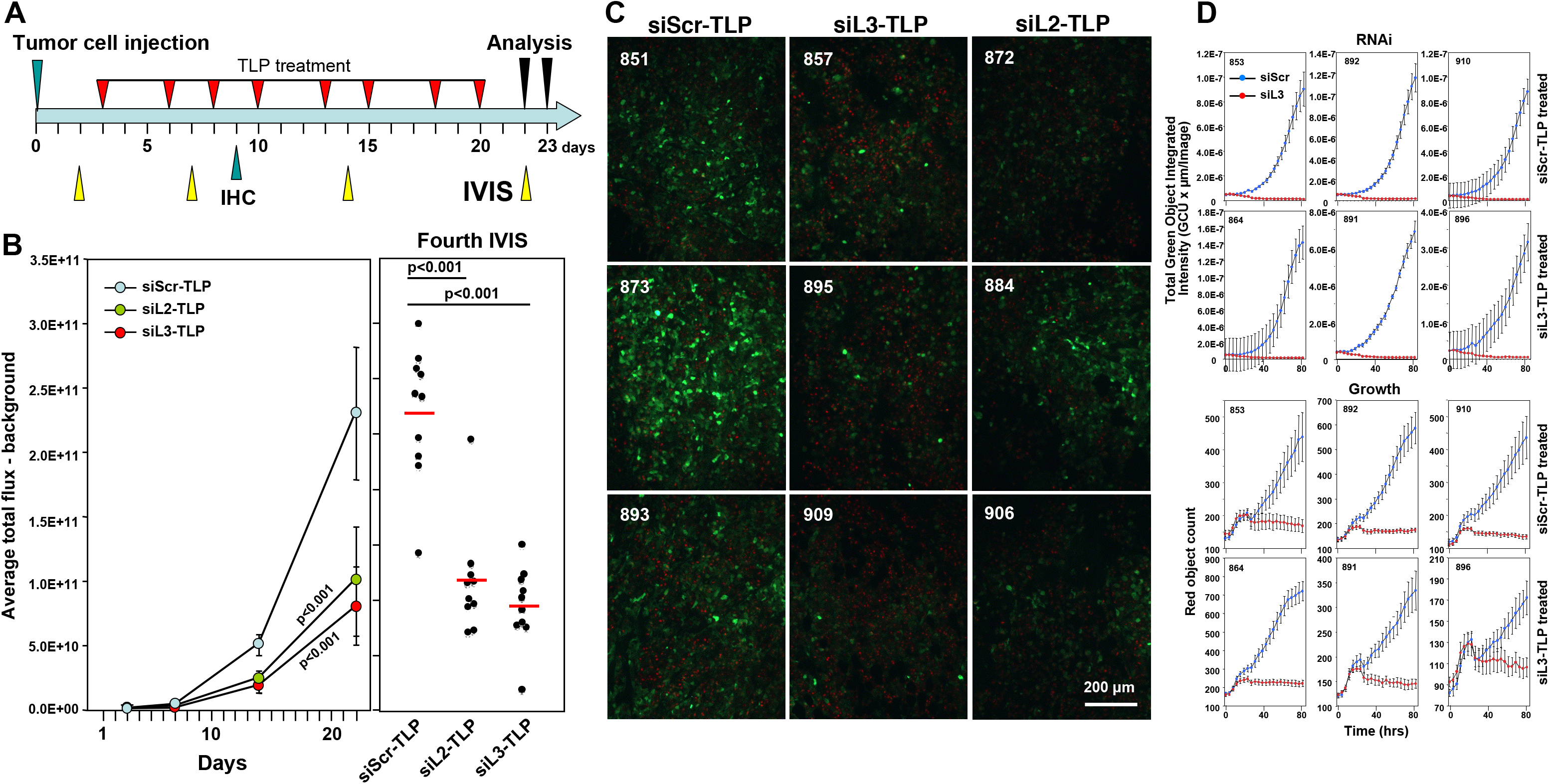
Tumors do not fully regress in response to siL3-TLP and siL2-TLP treatment but retain sensitivity to DISE. **A:** Treatment scheme. **B:** Small animal imaging of 100,000 HeyA8-Nuc-red-Luc-neo-Venus-CD95L cells injected i.p. into NGS mice treated with either siScr-TLPs, siL2-TLPs, or siL3-TLPs. *Left:* Tumor growth over time. *Right:* Bioluminescence signal in individual mice at the fourth IVIS (22 days after tumor injection) treated as indicated following the treatment protocol outlined in A. ANOVAs were performed for pairwise comparisons of average flux over time between siScr and siL3 treated and siScr and siL3 mice, respectively. **C:** Immunofluorescence analysis of frozen tumors from three mice of each treatment group in B. Red, nuclei of tumor cells; Green, Venus fluorescence (mouse numbers indicated in top left). **D:** Change in green fluorescence (RNAi, top 6 panels) and change in red object count (growth, bottom 6 panels) of tumor cells from 3 mice per siScr-TLP and siL3-TLP treatment group after transfection with either siScr or siL3. 1000 cells per well were plated (mouse numbers indicated in top left).

Similar to the experiments in which we expressed shRNAs in cancer cells *in vivo* to reduce tumor growth, treatment of the HeyA8 tumors with TLP nanoparticles also did not completely eliminate the cancer cells. To determine the mechanism of treatment resistance we isolated tumor cells from mice that had been treated with either siScr-TLP or siL3-TLP particles (three mice each) and after establishing tissue culture *in vitro* we introduced the siL3 siRNA into the cells by transfection (Figure 6D). In all cases, regardless of whether cells were derived from siScr-TLP or siL3-TLP treated mice, when siL3 was introduced it efficiently reduced the Venus fluorescence to zero (Figure 6D, top panels) and effectively killed the cells (Figure 6D, bottom panels). These data demonstrated that similar to the Dox treated HeyA8-pTIP-siL3 cells, the cells did not become resistant to the induction of DISE when they were isolated *ex vivo* and cultured under standard laboratory conditions. In summary, our data suggest that it is possible to induce DISE *in vivo*. Further work is needed to understand the mechanism of DISE induction, and to optimize strategies aimed at inducing DISE *in vivo*.

## DISCUSSION

Most targeted cancer therapy is based on small molecule inhibitors or biological reagents that target an oncogene often overexpressed in cancer cells. However, such targeted therapy has not resulted in a cure for the majority of cancer patients with advanced cancer. Most often patients develop resistance to the treatment, either because tumors in relapsing patients carry mutations in the targeted gene or the tumors upregulate factors that render them resistant to the therapy [14, 15]. Using more of the same therapy is futile and thus attempts are being made to find alternative targets for therapy or to re-sensitize cancer cells to the targeted therapy. In fact, it has been suggested that we will need to find “a radically different therapeutic modality” [16]. It has also been argued that rather than targeting individual genes, one needs to identify and disrupt networks of genes [17]. We are now providing such an approach, a radically different way of attacking cancer, that affects networks of survival genes.

We recently identified toxic RNAi active sequences in the genes of CD95 and CD95L [9]. We had reported earlier that introducing such sequences in the form of siRNAs or shRNAs induces a combination of multiple cell death pathways (we now call DISE) that cancer cells have a hard time developing resistance to [8]. Our new data on treating mice with nanoparticles by delivering two of these siRNAs *in vivo* now suggest that, in theory, it should be possible to induce this mechanism selectively in tumor cells *in vivo*. While we did detect resistance to treatment both when DISE was initiated in mice by expressing an inducible CD95L derived shRNA and when delivering CD95L derived siRNAs using nanoparticles, in both cases cells retained their intrinsic sensitivity to DISE. Tumor cell sensitivity was confirmed *ex vivo* as seemly resistant tumor cells treated both with lentiviral shRNAs or with commercially available siRNAs under standard culture conditions, both exhibited growth reduction upon reintroduction of treatment. Ultimately, further study is required to maximize the therapeutic efficacy of DISE. We chose ovarian cancer in an attempt to somewhat localize the nanoparticles to one tissue site in the animals. Indeed, when injected i.p., few nanoparticles could be detected on the liver of the animals but most of them were found to decorate or localize within the tumors (Supplementary Figure S4A and data not shown).

A key question remaining to be addressed is the issue of general toxicity, whether normal cells will die by DISE. While our siRNAs were derived from human CD95L, due to the mechanism through which DISE works it is very likely (and we now demonstrate for siL3) that the siRNAs will also kill mouse cells. In fact DISE was first seen in the mouse cell line CT26 stably expressing human CD95L when we overexpressed human CD95L derived shL3 [8]. Only later we discovered that shL3 very efficiently killed these mouse cells by targeting a network of critical survival genes in the mouse cells (data not shown). Recently, we reported data to suggest that normal cells are protected from DISE by the cellular miRNAs [9] which are known to be globally downregulated in human cancers when compared to their normal tissues [18]. However, the question whether DISE affects normal cells requires further research.

We showed that DISE preferentially affects cancer stem cells (CSCs) [10]. This selective sensitivity suggested that the DISE mechanism may have a physiological role in protecting stem cells from neoplastic transformation and that it can be used to target CSCs. The question arising as with every therapy directed at CSCs is whether it will affect somatic stem cells. This will have to be tested.

We propose that DISE will predominantly affect cancer cells making it unnecessary to specifically deliver the drugs to the cancer cells. In this case, the TLP nanoparticles used in our study are dependent on the expression of the SR-B1 scavenger receptor, a receptor expressed by numerous cancer types. Our data support that siRNA-TLP may be used to induce DISE in different types of cancer via local or systemic administration. Further, other targeted vesicles, such as exosomes, may be loaded with DISE-inducing siRNAs whereby systemic delivery may be accomplished. Finally, generating exosomes or nanoparticles with mixtures of toxic siRNAs derived from CD95/CD95L [9], or other genes that contain such sequences, may allow more potent targeting of cancer cells. While the siRNAs used in our study are toxic to cancer cells, recent work from our group suggests that the human genome is filled with genes that contain such toxic sequences allowing screening for the ones that are most toxic to cancer cells and least toxic to normal cells and then using aforementioned delivery strategies for *in vivo* therapy.

## MATERIALS AND METHODS

### Cell lines and tissue culture

Base media were supplemented with 10% heat-inactivated fetal bovine serum (FBS; Sigma-Aldrich) and 1% 100x penicillin/streptomycin and L-Glutamine (Mediatech Inc.). Adherent cells were dissociated with 0.25% (w/v) Trypsin-0.53 mM EDTA solution (Mediatech Inc.). Cells were cultured in an atmosphere of air, 95% and 5% carbon dioxide (CO_2_) at 37°C. For lentivirus production 293T cells were used. 293T cells were cultured in Dulbecco’s Modified Eagle’s Medium (DMEM). The following cell lines were used: breast cancer: MCF-7 (NCI60 cell panel), MDA-MB-231 (ATCC HTB-26), and HCC-70 (ATCC CRL-2315) were cultured in supplemented RPMI1640 Medium (Mediatech Inc.), SKBR-3 (ATCC HTB-30), in McCoy’s 5a Medium Modified (ATCC); Ovarian cancer: OVCAR-3, OVCAR-4, OVCAR-5, and OVCAR-8 (all Tumor Biology Core, Northwestern University) were cultured in RPMI1640; Caov-3 (ATCC HTB-75) in DMEM. HeyA8, Monty-1, and SKOV3IP1 were obtained from Dr. E. Lengyel, University of Chicago. HeyA8 was cultured in RPMI1640, Monty-1 and SKOV3IP1 in DMEM that was further supplemented with 1% non-essential amino Acids 100x (BioWhittaker), 1% Sodium Pyruvate 100 mM (BioWhittaker), and 1% MEM Vitamins 100x solution (Mediatech Inc.). Glioblastoma cell lines T98G (ATCC CRL1690) and U-87 MG (ATCC HTB-14) were cultured in Eagle’s Minimum Essential Medium (EMEM) (ATCC). The neuroblastoma cell line NB-7, the colorectal cancer cell line HCT116 (ATCC CCL-247), and JA3 (a subclone of Jurkat, an acute T-cell leukemia cell line, University of Heidelberg, Germany) were cultured in RPMI1640. The hepatocellular carcinoma cell line HepG2 (ATCC HB-80645) was cultured in EMEM (ATCC). ID8, a mouse ovarian cancer cell line, was cultured in DMEM supplemented with 4% FBS, and 10 mg/l Insulin, 5.5 mg/l Transferrin, 6.7 μg/ml Selenium (ITS, Mediatech, Inc., 1:10 diluted).

### Lentiviral infection

50,000 to 100,000 cells per 6-well were plated and infected the following day. When the lentivirus titer was known, the cells were infected at a multiplicity of infection (MOI) of 3 or 5 in the presence of 8 μg/ml polybrene, for 24 hours. Otherwise virus mix was produced by transfecting 293T cells. 3-4 million 293T cells were plated in a 10 cm dish in antibiotics free medium. Next day, in one tube 60 μl of Lipofectamine^®^ 2000 Transfection Reagent was mixed with 1ml Opti-MEM medium (Gibco), and this tube was incubated for 5 min at RT. In a second tube, a lentiviral vector, a packaging plasmid pCMVDR8.9, and an envelope plasmid pMD.G (VSV.G-envelope protein encoding plasmid) were mixed at a ratio of 12 μg:6 μg:6 μg in 1 ml Opti-MEM I. The two tubes were then mixed and incubated for 20 mins after which the transfection mix was added drop wise to the 10 cm dish of 293T cells. After 9 hours, the transfection mix was removed, and replaced with fresh full medium. The virus mix was collected 48 hours after transfection, sterile filtered, and aliquots were stored at −80°C until use. Cells were infected using a volume of 0.5 to 2 ml of this virus mix per 6-well. Where applicable, infected cells were selected with 3 μg/ml puromycin. To induce the expression of shRNAs from pTIP vectors the tetracycline analogue Doxycycline (Dox) was added to the culture at a final concentration of 0.1 μg/ml. To label nuclei red cells were infected with a puromycine selectible NucLight Red lentivirus (Essen Bioscience).

### Lentiviral vectors

To determine whether RNAi active sequences are taken up by the cells and active, cells were infected with lentivirus Venus sensors. They contain either a minisensor comprised of the 50 nucleotides surrounding the siL3 target site in CD95L (Venus-siL3WT), the mutated siL3 target site (Venus-siL3MUT) (described below), or the entire open reading frame of the CD95L-gene (NCBI accession number NM_000639.1) as recently described [9].

The Venus-siL3 sensor vector was created by subcloning an insert containing the Venus ORF followed by an artificial 3’UTR composed of a 62 bp portion of the CD95L cDNA containing the siL3 target site (5’-*GCCCTTCAATTACCCATAT*-3’) into a modified pCD510B vector (System Bioscience) as the backbone. IDT synthesized the insert as both a sense and antisense DNA strand containing an XbaI restriction enzyme (RE) site at the 5’ end and an EcoRI RE site at the 3’ end. The annealed insert and the modified pCD510B vector were digested with XbaI (NEB #R0145) and EcoRI (NEB #R0101). Subsequent ligation with T4 DNA ligase (NEB #M0202) created the pCD510B Venus-siL3 sensor vector. The Venus-siL3MUT sensor was generated by subcloning the Venus-siL3MUT insert into a modified CD510B-1 backbone [10] using XbaI and EcoRI. The insert was ordered as a synthetic gene from IDT containing an XbaI site and EcoRI site at the 5’ and 3’ ends, respectively and was composed of the Venus open reading frame followed by the sequence 5’-CTCGAGAGCTGCCGTGCAGCAGGACTTCAACTAGACATCTCCCCAGATCTACTGGG-3’, which contains the mutant siL3 sense sequence.

To monitor the growth of tumor cells in NSG mice by quantifying bioluminescence, the cells were infected with lentivirus pFU-Luc2-eGFP (pFUL2G), pFU-Luc2-tdTomato (pFUL2T) (a kind gift of Dr. Sanjiv Sam Gambhir at Stanford University, Stanford CA), or FUW-LucNeo (LucNeo) (G418 selectable) (Received from Dr. Jian-Jun Wei, Northwestern University). To express shRNAs derived from the CD95 or CD95L gene, cells were infected with the following MISSION Lentiviral Transduction Particles (the active sequence is underlined in each sh-RNA loop sequence): plko-shSCR = MISSION^®^pLKO.1-puro Control Non-Mammalian shRNA control Transduction articles (Sigma, SHC002V) as non targeting control (CCGG<u>CAACAAGATGAAGAGCACCAA</u>CTCGAGTTGGTGCTCTTCATCTTGTTGTTTTT). Plko-shL3: MISSION^®^ shRNA Lentiviral Transduction Particle for human CD95L (Sigma, Cat.No. SHCLNV-NM_000639 on exon 4, RCN0000059000) (CCGG<u>ACTGGGCTGTACTTTGTATAT</u>CTCGAGATATACAAAGTACAGCCCAGTTTTTTG). Plko-shR6: MISSION^®^ shRNA Lentiviral Transduction Particle for human CD95 (Sigma, Cat.No. SHCLNV-NM_000043 on exon 4, TRCN0000038696) (CCGG<u>GTGCAGATGTAAACCAAACTT</u>CTCGAGAAGTTTGGTTTACATCTGCACTTTTTG). These same shRNA sequences were cloned into the tetracycline inducible expression vector pTIP as previously described [8]. Cancer cells were infected with lentivirus for pTIP-shScr, or pTIP-shL3.

### Western blot analysis

Protein extracts were collected by lysing cells with RIPA lysis buffer [150 mM NaCl, 10 mM Tris HCl ph7.2, 1% SDS, 1% Triton X-100, 1% deoxycholic acid, 5 mM EDTA, Protein inhibitor cocktail tablet (1 tablet per 10 ml lysis buffer)]. 200 μM PMSF was added to the lysis buffer prior to use. Protein concentration was quantified using the DC^™^ Protein Assay (Bio-Rad). 30 μg of protein were resolved on 10% SDS-PAGE gels and transferred to nitrocellulose membranes (Amersham^™^ Protran ^™^, pre size 0.45 μm GE Healthcare Life Science) overnight at 25 mA. To verify the protein bands, a protein size marker was included on the gel, Amersham ECL Rainbow Molecular Weight Marker (GE Healtcare Life Science, Cat.No.: RPN800E). Membranes were incubated with blocking buffer (5% non-fat milk in PBST (PBS + 0.1% Tween-20) for 1 hour at room temperature. Membranes were then incubated with the primary antibody diluted in blocking buffer over night at 4°C. Membranes were washed 3 times with PBST. Secondary antibodies were diluted in blocking buffer and applied to membranes for 1 hour at room temperature. After 3 more additional washes, detection was performed using the ECL^™^ Western Blotting Detection Reagents reagent (GE Healthcare) or SuperSignal^™^ West Dura Extended Duration Substrate (ThermoFisher Sci.) and visualized with the chemiluminescence imager G:BOX Chemi XT4 (Syngene). All primary and secondary antibodies were diluted in a blocking buffer (5% non-fat milk in PBST) at different dilutions. The following primary antibodies were used: anti-human Fas antibody (C-20) Polyclonal rabbit IgG (Santa Cruz Biotechnology Inc., Cat.No.: sc-715, 1:500), anti-Scavenger receptor type B-1 rabbit IgG monoclonal [EP1556Y] (Abcam, Cat.No.: ab52629, 1:2000), and anti-Actin (I-19) goat polyclonal IgG (Santa Cruz Biotechnology Inc., Cat.No.: sc-1616, 1:2000). All secondary antibodies labeled with horse radish peroxidase: goat anti-rabbit IgG-HRP (Santa Cruz Biotechnology, Inc.; Cat.No.: sc-2004; 1:8000), rabbit anti goat IgG (Human adsorbed)-HRP (Santa Cruz Biotechnology Inc. sc-2768; 1:8000), goat-anti-rabbit-Ig human adsorbed-HRP (Southern Biotech; Cat.No.: 4010-05, 1:10000), or rabbit-anti-goat IgG(H+L) human adsorbed-HRP (Southern Biotech; Cat.No.: 6164-05, 1:10000).

### CRISPR/CAS9 genome editing to delete the entire CD95L gene in 293T cells

The procedure to generate specific mutant cells using two guide RNAs was recently described [9]. In short, the pMJ920 Cas9 vector (expressing a Cas9-GFP conjugate) was transfected using Lipofectamine^™^ 2000 (Invitrogen, Cat# 11668-019) with two guide RNA scaffolds containing guide RNA sequences targeting both 5’ and 3’ of the FASLG mRNA sequence to generate genomic human CD95L knock out cells. The following guide RNA were designed using the online CRISPR design tool (crispr.mit.edu described in [19]. Guide sequences with a quality score >70 were tested. The final guides used to generate FASLG knockouts are as follows (PAM sequence is underlined): 5’ guide sequence (TTGTGGGCGGAAACTTCCA<u>GGG</u>), 3’ guide sequence (GTACTGCCTATGTAAGCAC<u>TGG</u>), were transfected as G-blocks (IDT) into 293T cells. As a control, the Cas9 vector was transfected alone, without the addition of G-blocks encoding guide RNAs. The G block sequences were adopted from [20] and are as follows: 5’ Guide TGTACAAAAAAGCAGGCTTTAAAGGAACCAATTCAGTCGACTGGATCCGGTACCAAGGTCGGGCAGGAAGAGGGCCTATTTCCCATGATTCCTTCATATTTGCATATACGATA CAAGGCTGTTAGAGAGATAATTAGAATTAATTTGACTGTAAACACAAAGATATTAGTACAAAATACGTGACGTAGAAAGTAATAATTTCTTGGGTAGTTTGCAGTTTTAAAATTATGTTTTAAAATGGACTATCATATGCTTACCGTAACTTGAAAGTATTTCGATTTCTTG GCTTTATATATCTTGTGGAAAGGACGAAACACC**G<u>TTGTGGGCGGAAACTTCCA</u>**GTT TTAGAGCTAGAAATAGCAAGTTAAAATAAGGCTAGTCCGTTATCAACTTGAAAAAGTGGCACCGAGTCGGTGCTTTTTTTCTAGACCCAGCTTTCTTGTACAAAGTTGGCATTA 3’ Guide TGTACAAAAAAGCAGGCTTTAAAGGAACCAATTCAGTCGACTGGATCCGGTACCAAGGTCGGGCAGGAAGAGGGCCTATTTCCCATGATTCCTTCATATTTGCATATACGATA CAAGGCTGTTAGAGAGATAATTAGAATTAATTTGACTGTAAACACAAAGATATTAGTACAAAATACGTGACGTAGAAAGTAATAATTTCTTGGGTAGTTTGCAGTTTTAAAATTATGTTTTAAAATGGACTATCATATGCTTACCGTAACTTGAAAGTATTTCGATTTCTTG GCTTTATATATCTTGTGGAAAGGACGAAACACC**G**<u>TAATAGAGTGGCTTAGTAG</u>GTTT TAGAGCTAGAAATAGCAAGTTAAAATAAGGCTAGTCCGTTATCAACTTGAAAAAGTGGCACCGAGTCGGTGCTTTTTTTCTAGACCCAGCTTTCTTGTACAAAGTTGGCATTA. Cells successfully transfected were enriched by FACS sorting the top 20% GFP fluorescing cells three days after transfection (Day 1 transfection, Day 2 change media, Day 4 enrichment). The BD FACSAria SORP system was used. The cells were replated to recover. After 13 days in culture, single cells were sorted into 96 well plates containing conditioned media. Homozygous knockout clones were identified 3-4 weeks later by genomic PCR. The genotype of the 293T clones was determined by genomic PCR with external primers designed using Primer3: Lg5P_del_F (CATAAAATTATAGCCCCACTGACC) and Lg5P_del_R (CTGGGATGACAGCTTAAAGAAAAT), and internal primers FasLg_(int)_F (GTGGTAGGCTATTGTCCCTGGAAT) and FasLg_(int)_R (TGCAAGATTGACCCCGGAAGTATA) (IDT).

### Transfection with short oligonucleotides

For transfection of cancer cells with siRNAs Lipofectamine^®^ RNAiMAX or 2000 transfection reagent was used at a concentration that was optimized for each cell line, following the instructions of the vendor. The same sequences were ordered from two different vendors, Dharmacon and Integrated DNA Technologies (IDT): Dharmacon: Lyophilized ON-Target plus siRNA were resuspended in 1 x siRNA buffer (using 5x buffer from Dharmacon, through Thermo Fisher Scientific Biosciences, Cat. No.: B-002000-UB, diluted with RNAse/DNAse free water, to 60 mM KCl, 6 mM HEPES-pH 7.5, 0.2 mM MgCl_2_) to a concentration of 20 μM. IDT: Individual RNA oligos were ordered for the sense and antisense oligo; the sense strand had Ts added to the 3’ end; antisense strand had 2 deoxy As at the 3’ end, and phosphate residue on the 5’ end. For the visualization of siRNA-uptake a Cy5 label was attached to the 5’ end of the sense strand. Sense and antisense oligos were first resuspended in water at 500 μM (stock), then sense and antisense oligos were mixed with nuclease free Duplex buffer (IDT, Cat.No# 11-01-03-01; 100 mM Potassium Acetate, 30 mM HEPES, pH 7.5) to 20 μM (working solution), heated up for 2 minutes at 94°C, then the oligos were allowed to cool down to room temperature for 30 minutes. All siRNA solutions were aliquoted and stored at −80°C. The cells were transfected with siRNAs at a final concentration of 5 nM - 25 nM. The following siRNA sequences were used: siNT#2: UGGUUUACAUGUCGACUAA (Dharmacon D-001810-02, non targeting in mammalian cells), siL2: CAACGUAUCUGAGCUCUCU (Dharmacon J-011130-06, human CD95L exon 4), siL3: GCCCUUCAAUUACCCAUAU (Dharmacon J-011130-07, human CD95L exon 1.), siL3MUT: G<u>GA</u>CUUCAA<u>C</u>UA<u>GA</u>CAU<u>C</u>U (siL3 sequence with 6 changes), siL3-Cy5: GCCCUUCAAUUACCCAUAU-Cy5 (siL3 sequence with Cy5 fluorophore).

### Total RNA isolation and RNA-seq analysis

HeyA8 cells were transfected in 6-wells with IDT siNT2 or siL3 oligonucleotides at 25 nM. The transfection mix was removed after 9 hours. Total RNA was isolated 48 hours after initial transfection using the miRNeasy Mini Kit (Qiagen, Cat.No. 74004)) following the manufacturers instructions. An on column digestion step using the RNAse-free DNAse Set (Qiagen, Cat.No.: 79254) was included. NGS RNA-SEQ library making and sequencing was performed by the University of Chicago Genomics Facility. The quality and quantity of RNA samples was assessed using an Agilent bio-analyzer. RNA-SEQ libraries were generated using Illumina Stranded TotalRNA TruSeq kits using the Illumina provided protocol and sequencing was performed using the Illumina HiSEQ4000 using Illumina provided protocols and reagents. Sequences were aligned to the human genome and analyzed as recently described [9]. The accession number for the RNA-Seq and expression data reported in this paper is GSE101167.

### Monitoring growth and fluorescence expression over time

To monitor cell growth over time, cells were seeded between 125 or 10,000 per well in a 96-well plate in triplicates. The plate was then scanned using the IncuCyte ZOOM live cell imaging system (Essen BioScience). Images were captured at regular intervals, at the indicated time points, using an 10x objective. Cell confluence, red object count, and the green object integrated intensity were calculated using the IncuCyte ZOOM software (version 2015A).

### Synthesis of templated lipoprotein particles (TLP) and siRNA-TLPs

For TLP synthesis, an aqueous solution of citrate stabilized gold nanoparticles (Au NP) (80 nM, 5 ± 0.75 nm, Ted Pella, Inc.) was mixed with a 5-fold molar excess of purified human apolipoprotein A-I (apoA-I) (400 nM, Meridian Life Sciences, >95% pure by SDS PAGE) in a glass vial. The AuNP/ apo A-I mixture was incubated overnight at room temperature (RT) in a flat bottom shaker at low speed. Next, a 1:1 ratio of two phospholipids: 1,2-dipalmitoyl-sn-glycero-3-phosphoethanolamine-N-[3-(2-pyridyldithio) propionate] (PDP-PE) and 1,2-dioleoyl-sn-glycero-3-phophocholine (DOPC) (Avanti Polar Lipids), each dissolved in chloroform (CHCl3, 1 mM), were added to the AuNP/ apo A-I solution in 250-fold molar excess to the AuNP. PDP-PE was added first and the solution was vortexed prior to adding DOPC. Next, cholesterol dissolved in CHCl_3_ (1 mM, Sigma Aldrich) was added in 25-fold molar excess to the AuNP. The mixture was vortexed and briefly sonicated (~ 2 min) causing the solution to become opaque and pink in color. The resulting mixture was gradually heated to ~65°C with constant stirring to evaporate CHCl3 and to transfer the phospholipids onto the particle surface and into the aqueous phase (~20 minutes). The reaction was complete when the solution returned to a transparent red color. The resultant TLPs were incubated overnight at RT and then purified via centrifugation (15,870 x g, 50 min) or tangential flow filtration (TFF). The supernatant was removed in the case of centrifugation and the resulting purified and concentrated TLPs were combined into a single vial. TLPs were stored at 4°C until use. The concentration of the TLPs was measured using UV-Vis spectroscopy (Agilent 8453) where AuNPs have a characteristic absorption at λ_max_ = 520 nm, and the extinction coefficient for 5 nm AuNPs is 9.696 x 10^6^ M^-1^cm^-1^.

To synthesize siRNA-TLP, RNA and 1,2-dioleoyl-3-trimethylammonium-propane (DOTAP) were first mixed. Individual sense and antisense RNA sequences of the siL2, siL3, or scrambled (siScr) (IDT) were re-suspended in nuclease free water (500 μM, final). Complement pairs were then mixed in nuclease free water at a concentration enabling direct addition to TLPs (100 nM) at 25-fold molar excess of each RNA sequence (2.5 μM, final per RNA sequence). An ethanolic (EtOH) solution of DOTAP was then added to the RNA mixture to desired DOTAP:RNA molar ratios. In each case the resulting solvent ratio was 9:1, EtOH:water (v/v). The mixture of DOTAP and RNA was briefly sonicated and vortexed (x3) and then incubated at RT for 15 minutes prior to addition to a solution of TLPs in water. After the DOTAP-RNA mixture was added to the TLPs, the solvent mixture was 9:1, water:EtOH (v/v). The solution was incubated overnight at RT with gentle shaking on a flat bottom shaker at low speed. Resulting siRNA-TLPs were purified via centrifugation (15,870 x g, 50 min), the supernatant with unbound starting materials was removed, the pellets were briefly sonicated and then combined in a single tube to concentrate the siRNA-TLPs. The concentration of the siRNA-TLPs was calculated as described for TLP. For siRNA-TLPs, a strong absorption at λm_ax_ = 260 nm confirmed the presence of RNA.

### Monitoring of Cy5-labeled particle uptake

For FACS analysis, cells were plated at 50,000 cells per 12-well plate in 1 ml medium. After about 16 hours the entire medium was replaced with medium containing 10 nM TLP. After 24 hours incubation with the particles, the cells were harvested, trypsinized when necessary, and analyzed with FACS. For IncuCyte monitoring, cells were plated at 10000, 5000, 2500, and 1250 cells per 96-well. Images for phase, and the red channel were taken every 2.5 hours.

### Intra peritoneal (*i.p.*) injection of ovarian cancer cells into NSG mice and *in vivo* imaging

10^5^-10^6^ modified HeyA8 (always infected with a luciferase lentivirus) were injected i.p. into 6-week-old female NSG mice (NOD *scid* gamma, NOD-scid IL2Rg^null^, NOD-scid IL2Rgamma^nul^) (Jackson Laboratory, Cat.No.: 005557) following the Northwestern University Institutional Animal Care and Use Committee (IACUC)-approved protocol. The growth of tumor cells in the mice over time was monitored non-invasively using the IVIS^®^ Spectrum *in vivo* imaging system (Perkin Elmer). The tumor load was quantified by the luminescence of the regions of interest (the same area for each mouse encompassing the entire abdomen) using the Living Image software. To visualize the tumor load in the mice, 200 μl of the sterile Luciferin stock solution (15 mg/ml in Dulbeccos’s phosphate buffered Saline without Mg^2+^ and Ca^2+)^ was i.p. injected per mouse (which equals approximately 10 μl of luciferin stock per g of body weight). The mice were imaged 15 minutes after injection. Total Flux values of the different treatment groups were compared.

### H&E staining

Tissue samples or tumors were fixed in 10% Normal Buffered Formalin (VWR, Cat.No.: 16004-128) for 16 hrs, and further processed by the Northwestern University Mouse Histology & Phenotyping Laboratory (MHPL) (paraffin embedding, sectioning at 4 μM, slide preparation, and staining). Histology services were provided by the Northwestern University Research Histology and Phenotyping Laboratory at the Robert H Lurie Comprehensive Cancer Center.

### Microscopy and imaging

The following microscopes were used for imaging: Zeiss Axioscope with Nuance camera (Nuance multispectral imaging system) with the software called Nuance 3.0.0. to image Venus green and tdTomato Red. Imaging work on this microscope was performed at the Northwestern University Center for Advanced Microscopy. A Zeiss Axiovert S100 fluorescence microscope equipped with an Axiocam digital camera, and the software AxioVision Rel 4.8. was used to image samples with Venus green and Red Fluorescent protein. Images of histological slides, H&E images were taken on a Leica DM 4000B Light microscope D (Leica Microsystems) equipped with a Leica DFC320 color digital camera (Leica Microsystems). The Software program used to capture the images was called Leica Application Suite version 44.0 (Build:454).

### RNAi targeting of Venus-sensor - analysis by FACS

To analyze tumors by FACS, tumors were isolated from the peritoneal cavity of a mouse, washed in PBS, and dissociated with Trypsin. After the cell suspension was strained over a 70 μm Fisherbrand^®^ nylon mesh (Fisher Scientific, Cat.No.: 22363548), the suspension over the cells were analyzed by FACS. BD LSRFortessa Analyser at the Flow Cytometry Core, Northwestern University, was used for flow analyses. The data was analyzed using the program FlowJo 8.8.6. To ensure that only tumor cells were analyzed, only the red events (NucRed or pFUL2T infected cancer cells) were considered. The percentage of high green and low green expressing cells were quantified. For assessing the granularity of the tumors cells, the values of the side scatter area were compared between treatments.

### RNAi targeting of Venus-sensor - analysis by fluorescent microscopy

Tumors were fixed in 4% Paraformaldehyde (A 32% Paraformaldehyde stock solution, EM grade (Electron Microcopy Sciences, Cat.No.: 15714-S) was diluted with deionized water) over night, followed by a sucrose gradient (10 % sucrose for 2 hours, 20 % sucrose for 2 hours, 30 % sucrose overnight), before they were shock frozen in Tissue-Plus^®^ O.C.T. Compound Embedding medium (Scigen Scientific Gardena, CA, Cat.No.: 4583) following standard tissue freezing procedures. The frozen O.C.T blocks were sectioned at 6 um, mounted on slides by the MHPL (Northwestern University), and stored at −80°C until use. To assess the effect of the treatment on the Venus-fluorescence, slides with the section was thawed at room temperature, washed gently three times with PBS buffer, and mounted with a coverslip using Antifade VectaShield + DAPI (Vector Laboratories Inc., CA, Cat.No.: H-1200). Images of tumors with different treatments were taken on the Zeiss Axiovert S100 fluorescence microscope with the identical settings. Tumor cells only expressed a red fluorophore, which allowed to compare tumor areas with high as well as low green fluorescent Venus-sensor expression.

### Serum collection and toxicity test

Two mice from each TLP treatment group were selected as representatives to assess TLP toxicity. Whole blood was collected from each mouse via retro-orbital bleeding using disposable glass Pasteur pipettes when mice were sedated. Blood was then allowed to gravity drip into Greiner Bio-One^™^ MiniCollect^™^ Capillary Blood Collection System Tubes (Fisher Scientific, Cat.No.: 22-030-400) that were kept on ice. The collection tubes containing the serum were spun at 3000 g for 10 minutes at 15° C. After completion of the spin, two layers formed in the tube. The top layer containing the whole serum was collected, volume measured, transferred to new sterile Eppendorf^™^ Snap-Cap Microcentrifuge Safe-Lock^™^ Tubes (Fisher Scientific, Cat.No.: 05-402-25), frozen, and stored at −20° C. The serum samples were analyzed using a Complete Chemistry Profile Test performed by the Charles River Research Animal Diagnostic Services (Wilmington, MA).

### Statistical methods

Two-way analysis of variances (ANOVA) were performed using the STATA14 software to compare growth curves. One-tail student t-test was performed in the software package R to compare tumor load between treatment groups. Wilcoxon Rank Sum test was performed in R to compare IVIS signal between treatment groups.

## ABBREVIATIONS

DISE: Death induced by survival gene elimination
TLP: templated lipoprotein particles
UTR: untranslated region
OTE: off-target effect
MOI: multiplicity of infection
Dox: doxycycline
nt: nucleotide
RNAi: RNA interference

## AUTHOR CONTRIBUTIONS

A.E.M., A.H.K., M.P., N.R., C.L., S.B. performed experiments. K.M. and C.S.T. generated the nanoparticles, J.J.W. provided pathology expertise and M.E.P. directed the study and wrote the manuscript.

## ACKNOWLEDGEMENT

We are grateful to Dr. Stijn van Dongen for performing the Sylamer analysis and to William Putzbach for performing the GSEA.

---

**Supplementary Figure S1:**
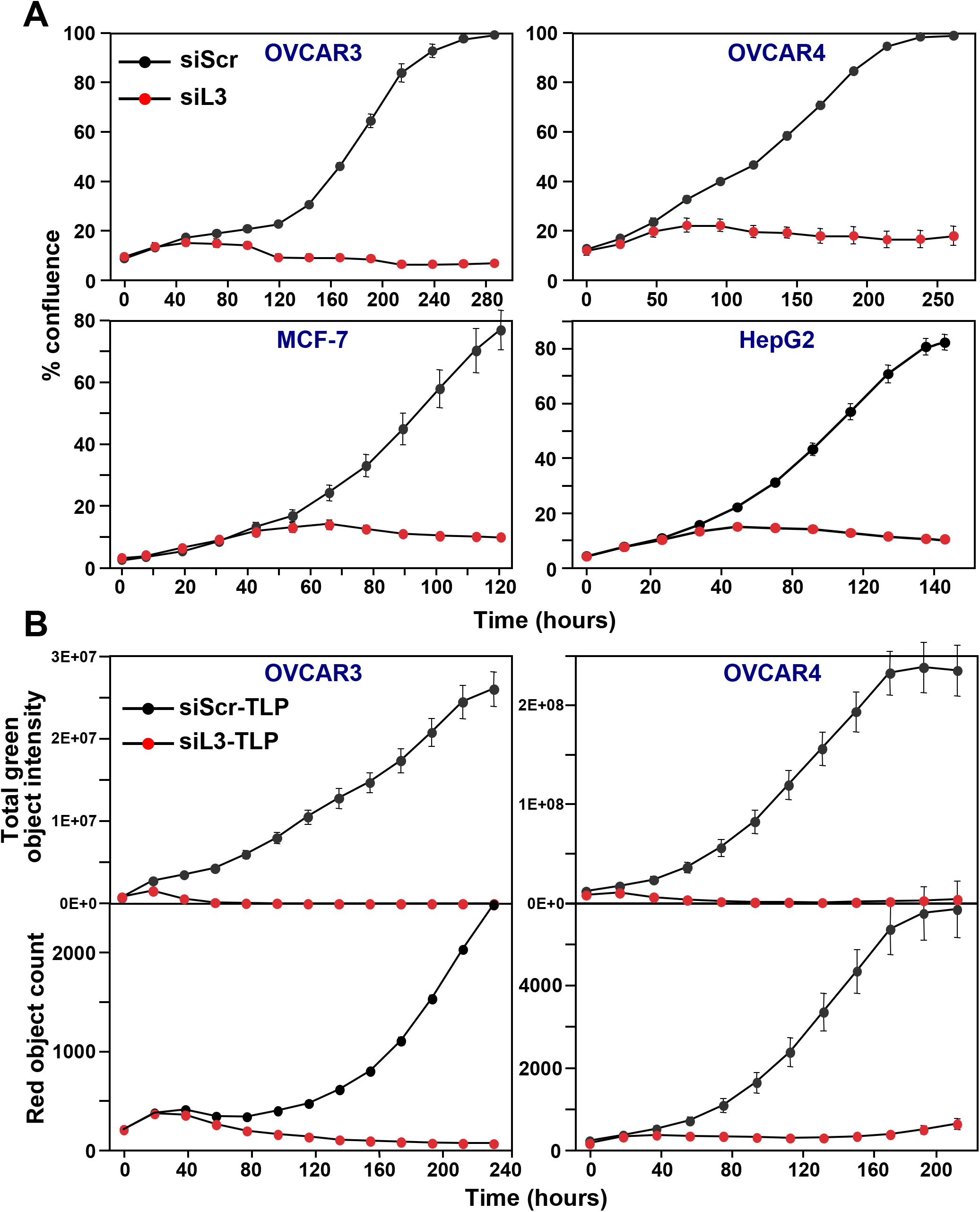
Induction of DISE by siL3 in different cancer cell lines. **A:** Confluency over time of four cancer cell lines transfected with 10 nM (OVCAR3, OVCAR4 or HepG2) or 25 nM (MCF-7) siScr or siL3. **B:** *Top:* Total green intensity or *bottom:* confluency over time of OVCAR3 Venus siL3 or OVCAR4 Venus siL3 cells treated with 30 nM of either siScr-TLP or siL3-TLP.

**Supplementary Figure S2:**
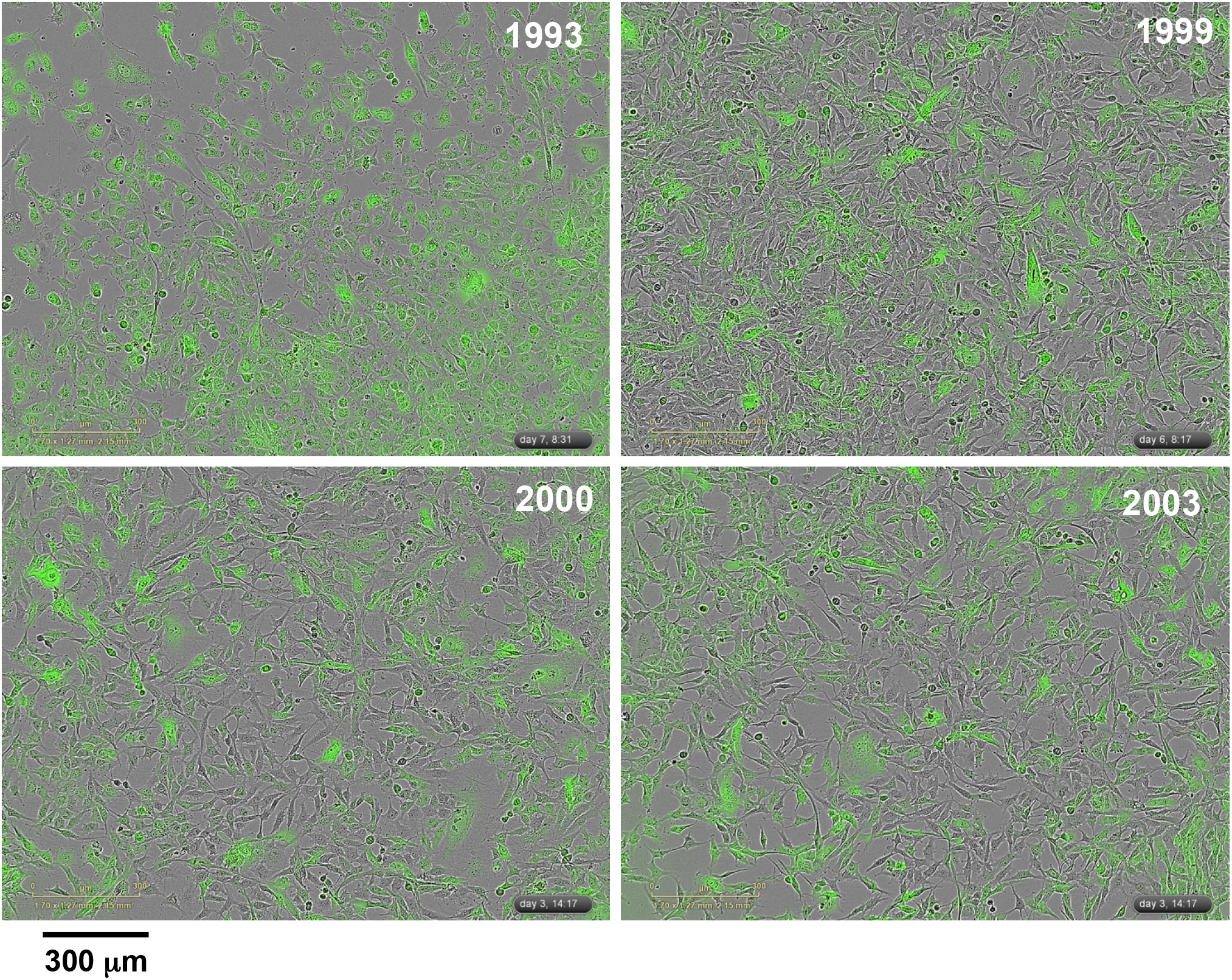
Combined phase contrast and green fluorescence of cells isolated from the four tumors used in Figure 2C.

**Supplementary Figure S3:**
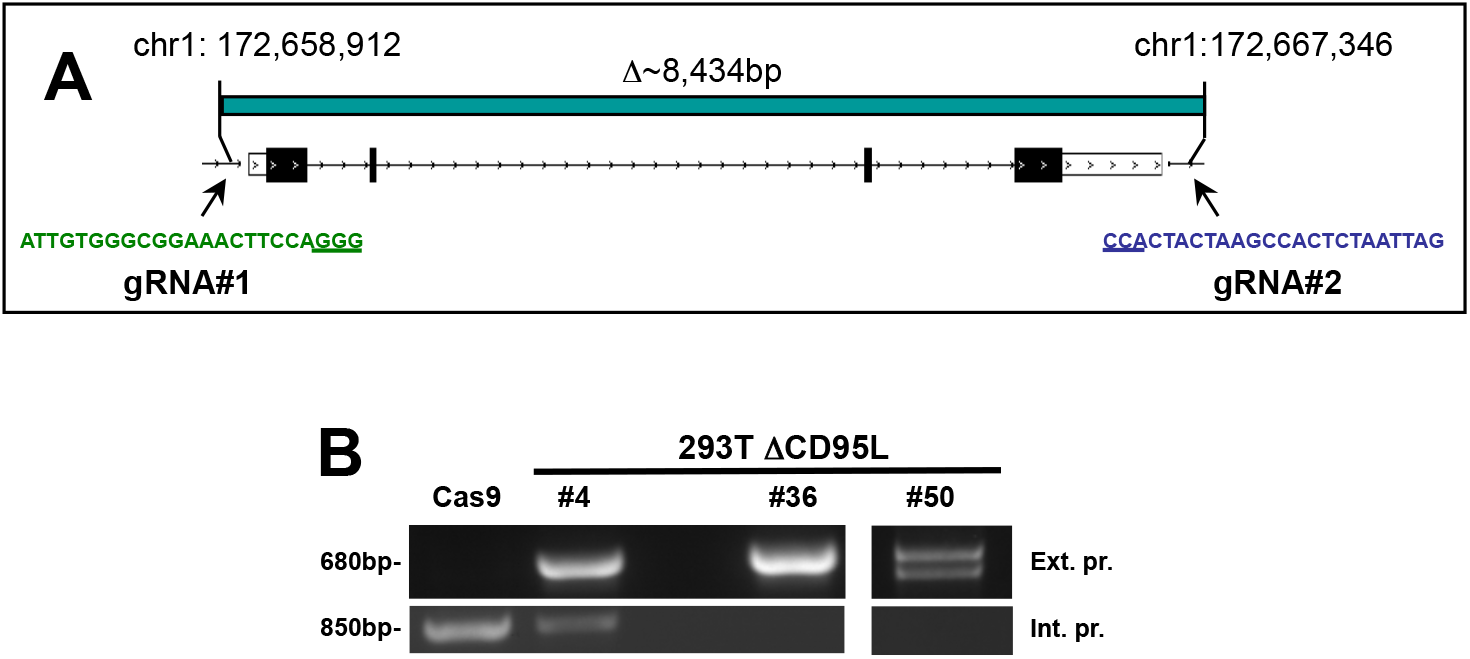
Generation of homozygous CD95L deletion in 293T cells. **A:** Schematic of the genomic locations and sequences of the gRNAs used to excise the entire CD95L gene. Blue indicates a gRNA targeting the antisense strand. **B:** PCR with flanking *(top panels)* and internal *(bottom panels)* primers was used to confirm the absence of CD95L in 293T clones. Cells infected with Cas9 only (Cas9) are wild-type, clone #4 is heterozygous, and clones #36 and #50 carry homozygous deletions.

**Figure S4:**
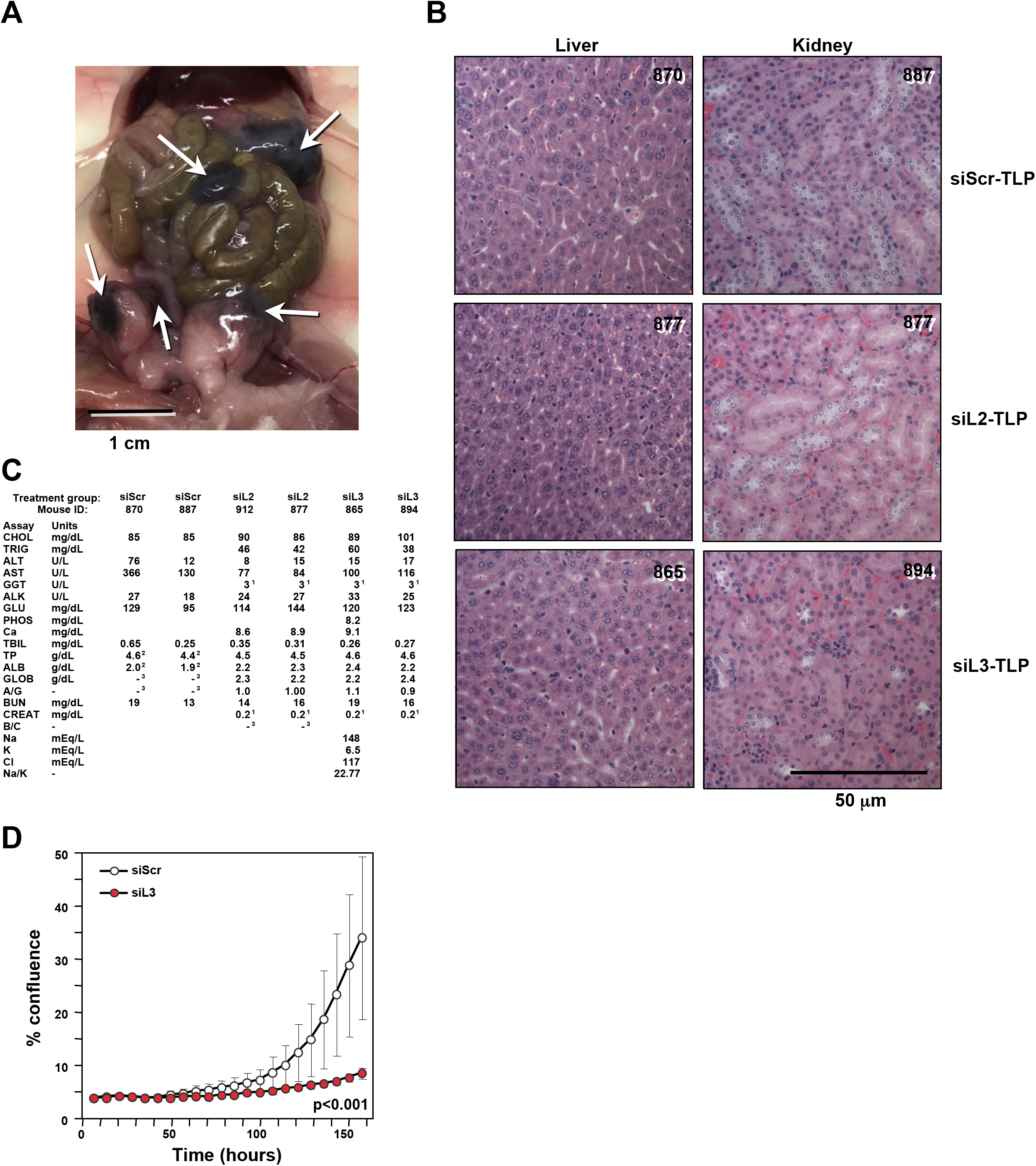
Supplementary No toxicity of siL3-TLP in NGS mice. **A:** Representative image of mouse peritoneum with HeyA8 tumor stained grey caused by the gold containing TLPs (arrows). **B:** Representative H&E staining of liver and kidney sections from mice of the three treatment groups. **C:** Serum analysis of two mice per treatment group. 1 = Sample assay value is less than the dynamic range. For most assays, the dynamic range low limit is reported. 2 = Sample was diluted for testing. Assay value for sample was below dynamic range, but results have been corrected for dilution. 3 = Assay is a calculated value. Either or both assay values used in the calculation were below the dynamic range of the assay, therefore no result is reported. **D:** Induction of DISE in ID8-Venus-mFLg cells after transfection with 25 nM siL3. As a control cells were transfected with siScr. ANOVA was performed for pairwise comparison of change in confluence over time between siScr and siL3 treated cells.

Movie S1: HeyA8-pTIP-shL3 cells.

Movie S2: HeyA8-pTIP-shL3 cells plus Dox.

Movie S3: HeyA8-pTIP-shL3 H5 clone growing in the presence of Dox. Resistant cells are growing into the field from the right.

Movie S4: HeyA8-pTIP-shScr control cells growing in the presence of Dox.

Movie S5: HeyA8 Venus siL3WT cells treated with siL3MUT.

Movie S6: HeyA8 Venus siL3WT treated with siL3.

Movie S7: HeyA8 Venus siL3MUT treated with siL3MUT.

Movie S8: HeyA8 Venus siL3MUT treated with siL3.

